# Insights into the genomic evolution of insects from cricket genomes

**DOI:** 10.1101/2020.07.07.191841

**Authors:** Guillem Ylla, Taro Nakamura, Takehiko Itoh, Rei Kajitani, Atsushi Toyoda, Sayuri Tomonari, Tetsuya Bando, Yoshiyasu Ishimaru, Takahito Watanabe, Masao Fuketa, Yuji Matsuoka, Austen A. Barnett, Sumihare Noji, Taro Mito, Cassandra G. Extavour

## Abstract

Most of our knowledge of insect genomes comes from Holometabolous species, which undergo the complete metamorphosis and have genomes under 2Gb with little signs of DNA methylation. In contrast, Hemiemetabolous insects undergo the ancestral incomplete metamorphosis and have larger genomes with high levels of DNA methylation. Hemimetabolous species from the Orthopteran order (grasshoppers and crickets) have some of the largest insect genomes. What drives the evolution of these unusual insect genome sizes, remains unknown. Here we report the sequencing, assembly and annotation of the 1.66-Gb genome of the Mediterranean field cricket *Gryllus bimaculatus*, and the annotation of the 1.60-Gb genome of the Hawaiian cricket *Laupala kohalensis.* We compare these two cricket genomes with those of 14 additional insects, and find evidence that hemimetabolous genomes expanded due to transposable element activity. Based on the ratio of observed to expected CpG sites, we find higher conservation and stronger purifying selection of methylated genes than non-methylated genes. Finally, our analysis suggests an expansion of the *pickpocket* class V gene family in crickets, which we speculate might play a role in the evolution of cricket courtship, including their characteristic chirping.

## Introduction

Much of what we know about insect genome structure and evolution comes from examination of the genomes of insects belonging to a single clade, the Holometabola. This group includes species such as flies and beetles, and is characterized by undergoing complete, or holometabolous, metamorphosis, in which the product of embryogenesis is a larva, which then undergoes an immobile stage called a pupa or chrysalis, during which the larval body plan is abandoned and the new, adult body plan is established. Following the pupal stage, the adult winged insect emerges^1^. This clade of insects includes nearly 90% of extant described insect species^2^. Members of this clade have become prominent model organisms for laboratory research, including the genetic model *Drosophila melanogaster.* Thus, a large proportion of our knowledge of insect biology, genetics, development, and evolution is based on studies of this clade.

Before the evolution of holometabolous metamorphosis, insects developed through incomplete or hemimetabolous metamorphosis. This mode of development is characterized by a generation of the final adult body plan during embryogenesis, followed by gradual physical growth of the hatchling through nymphal stages until the last transition to the sexually mature, winged adult, without major changes in body plan from hatchling to adult^1^. Many extant species maintained this presumed ancestral type of metamorphosis, including crickets, cockroaches, and aphids. Among hemimetabolous insects, most of our current genomic data is from the order Hemiptera (true bugs), which is the sister group to the Holometabola. For the remaining 15 hemimetabolous orders, genomic data remain scarce.

Based on data available to date, genome size and genome methylation show unexplained variation across insects. While most holometabolan species have relatively small genomes (0.2-1.5 pg), hemimetabolous species, and specifically polyneopterans (a taxon comprising 10 major hemimetabolous orders of winged insects with fan-like extensions of the hind wings), display a much larger range of genome sizes (up to 8 pg)^3^. This has led to the hypothesis that there is a genome size threshold at 2 pg (~2 Gb) for holometabolan insect genomes^3^. Studying genome size evolution the polyneopterans order Orthoptera (crickets, grasshoppers, locusts, and katydids) offers a valuable opportunity to investigate potential mechanisms of genome size evolution, as it includes species that have similar predicted gene counts, but have genomes ranging from 1.25 Gb to 16.56 Gb^4^. With respect to the level of CpG DNA methylation, only a few holometabolous species display evidence of genome wide DNA methylation at CpG sites, whereas 30 out of 34 studied polyneopteran species do^5,6^. However, the role of DNA methylation in polyneopteran species, and why it appears to have been lost in many holometabolans, is not clear.

Here, we present the 1.66-Gb genome assembly and annotation of *G. bimaculatus* (Orthoptera), commonly known as the two-spotted cricket, a name derived from the two yellow spots found on the base of the forewings of this species (**Figure 1A**). We also report the first genome annotation for a second cricket species, the Hawaiian cricket *Laupala kohalensis*, whose genome assembly was recently made public^7^. *G. bimaculatus* has been widely used as a laboratory research model for decades, in scientific fields including neurobiology and neuroethology^8,9^, evo-devo^10^, developmental biology^11^, and regeneration^12^. Technical advantages of this cricket species as a research model include the fact that *G. bimaculatus* does not require cold temperatures or diapause to complete its life cycle, it is easy to rear in laboratories since it can be fed with generic insect or other pet foods, it is amenable to RNA interference (RNAi) and targeted genome editing^13^, stable germline transgenic lines can be established^14^, and it has an extensive list of available experimental protocols ranging from behavioral to functional genetic analyses^15^.

**Figure 1:**
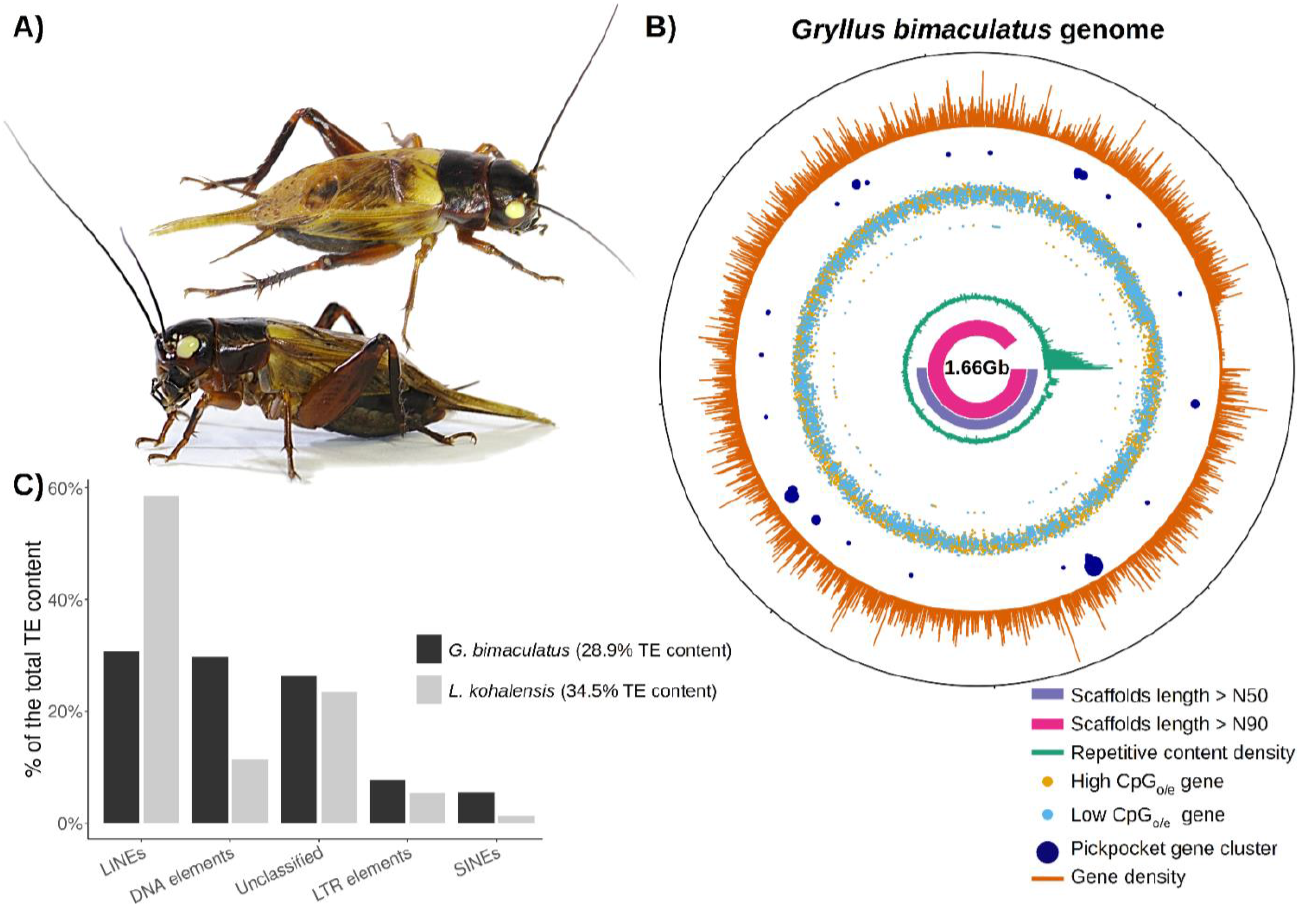
The *G. bimaculatus* genome. **A)** The cricket *G. bimaculatus* (top and side views of an adult male), commonly called the two-spotted cricket, owes its name to the two yellow spots on the base of the forewings. **B)** Circular representation of the *G. bimaculatus* genome, displaying the N50 (pink) and N90 (purple) scaffolds, repetitive content density (green), the high-(yellow) and low-(light blue) CpG_o/e_ value genes, *pickpocket* gene clusters (dark blue), and gene density (orange). **C)** The proportion of the genome made up of transposable elements (TEs) is similar between *G. bimaculatus* and *L. kohalensis* (28.9% and 34.5% respectively), but the specific TE family composition varies widely.

Comparing the two cricket genomes annotated here, with those of 14 other insect species, allowed us to identify three interesting features of these cricket genomes, some of which may relate to their unique biology. First, the differential transposable element (TE) composition between the two cricket species suggests abundant TE activity since they diverged from a last common ancestor, which our results suggest occurred circa 89.2 million years ago (Mya). Second, based on gene CpG depletion, an indirect but robust method to identify typically methylated genes^5,16^, we find higher conservation of typically methylated genes than of non-methylated genes. Finally, our gene family expansion analysis reveals an expansion of the *pickpocket* class V gene family in the lineage leading to crickets, which we speculate might play a relevant role in cricket courtship behavior, including their characteristic chirping.

## Results

### *Gryllus bimaculatus* genome assembly

We sequenced, assembled, and annotated the 1.66-Gb haploid genome of the white eyed mutant strain^12^ of the cricket *G. bimaculatus* **(Figure 1A)**. 50% of the genome is contained within the 71 longest scaffolds (L50), the shortest of them having a length of 6.3 Mb (N50), and 90% of the genome is contained within 307 scaffolds (L90). In comparison to other polyneopteran genomes, our assembly displays high quality in terms of contiguity (N50 and L50), and completeness (BUSCO scores) (**Supplementary Table 1)**. Notably, the complete BUSCO scores^17^ of this genome assembly at the arthropod and insect levels are 98.50% (C:98.5% [S:97.2%, D:1.3%], F:0.4%, M:1.1%, n:1066) and 97.00% (C:97.0% [S:95.2%, D:1.8%], F:0.8%, M:2.2%, n:1658) respectively, indicating high completeness of this genome assembly **(Table 1)**. The low percentage of duplicated BUSCO genes (1.31%-1.81%) suggests that putative artifactual genomic duplication due to mis-assembly of heterozygotic regions is unlikely.

**Table 1:** Gryllus bimaculatus *genome assembly statistics.*

|  |  |
| --- | --- |
| Number of Scaffolds | 47,877 |
| Genome Length (nt) | 1,658,007,496 |
| Genome Length (Gb) | 1.66 |
| Avg. scaffold size (Kb) | 34.63 |
| N50 (Mb) | 6.29 |
| N90 (Mb) | 1.04 |
| L50 | 71 |
| L90 | 307 |
| Complete BUSCO Score – Arthropoda | 98.50% |
| Complete BUSCO Score – Insecta | 97.00% |

### Annotation of two cricket genomes

The publicly available 1.6-Gb genome assembly of the Hawaiian cricket *L. kohalensis^7^*, although having lower assembly quality scores (N50=0.58 Mb, L90 = 3,483) than that of *G. bimaculatus*, scores high in terms of completeness, with BUSCO scores of 99.3% at the arthropod level and 97.80% at the insect level **(Supplementary Table 1)**.

Using three iterations of the MAKER2 pipeline^18^, in which we combined *ab-initio* and evidence-based gene models, we annotated the protein-coding genes in both cricket genomes (**Supplementary Figures 1 & 2**). We identified 17,871 coding genes and 28,529 predicted transcripts for *G. bimaculatus*, and 12,767 coding genes and 13,078 transcripts for *L. kohalensis* **(Table 2)**.

**Figure 2:**
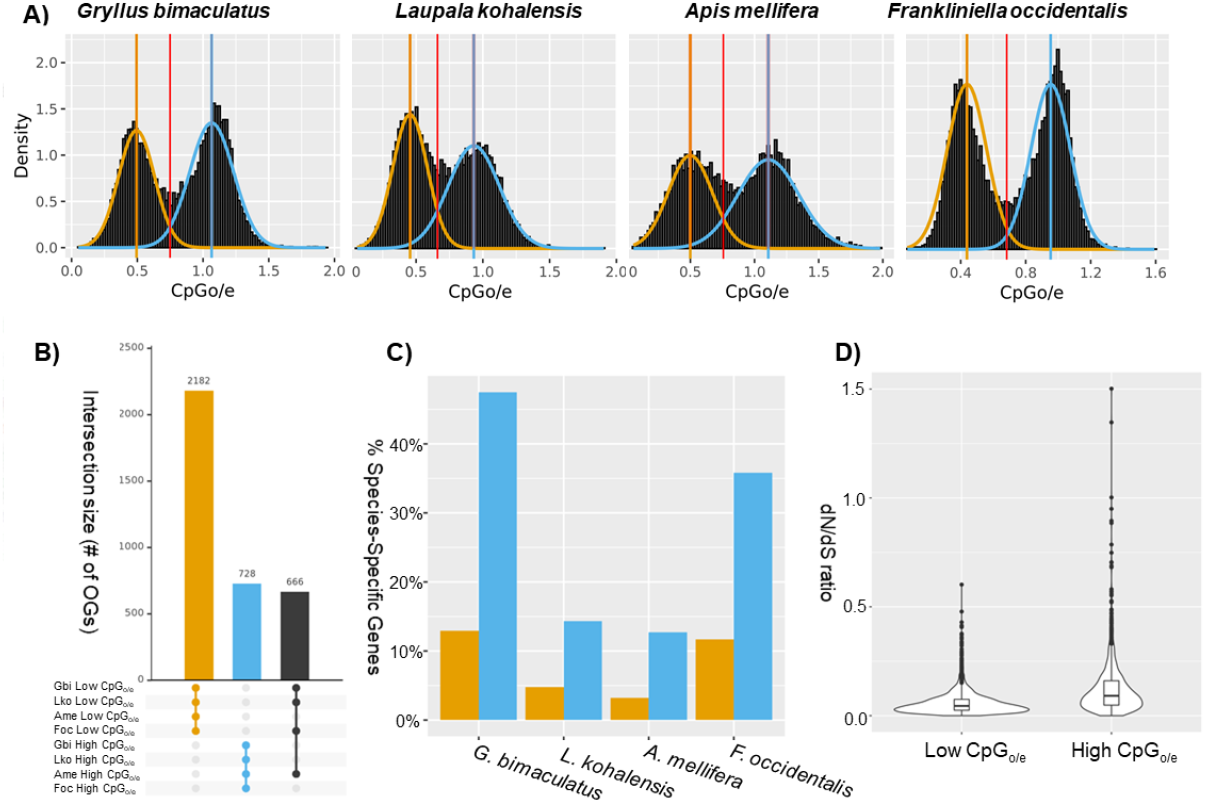
CpG_o/e_ distribution across insects. **A)** The distribution of CpG_o/e_ values within the CDS regions displays a bimodal distribution in the two crickets, as well as in the honeybee *A. mellifera* and the thrips *F. occidentalis*. We modeled each peak with a normal distribution and defined their intersection (red line) as a threshold to separate genes into low- and high-CpG_o/e_ value categories represented in yellow and blue color respectively. **B)** UpSet plot showing the top three intersections (linked dots) in terms of the number of orthogroups (OGs) commonly present in the same category (low-and high-CpG_o/e_) across the four insect species. The largest intersection corresponds to 2,182 OGs whose genes have low CpG_o/e_ in the four insect species, followed by the 728 OGs whose genes have high CpG_o/e_ levels in all four species, and 666 OGs whose genes have low CpG_o/e_ in the three hemimetabolous species and high CpG_o/e_ in the holometabolous honeybee. Extended plot with 50 intersections is shown in **Supplementary Figure 4**. **C)** Percentage of species-specific genes within low CpG_o/e_ (yellow) and high CpG_o/e_.(blue) in each insect, indicating that more such genes tend to have high CpG_o/e_ values. **D)** One-to-one orthologous genes with low CpG_o/e_ values in both crickets have significantly lower dN/dS values than genes with high CpG_o/e_ values.

**Table 2:**
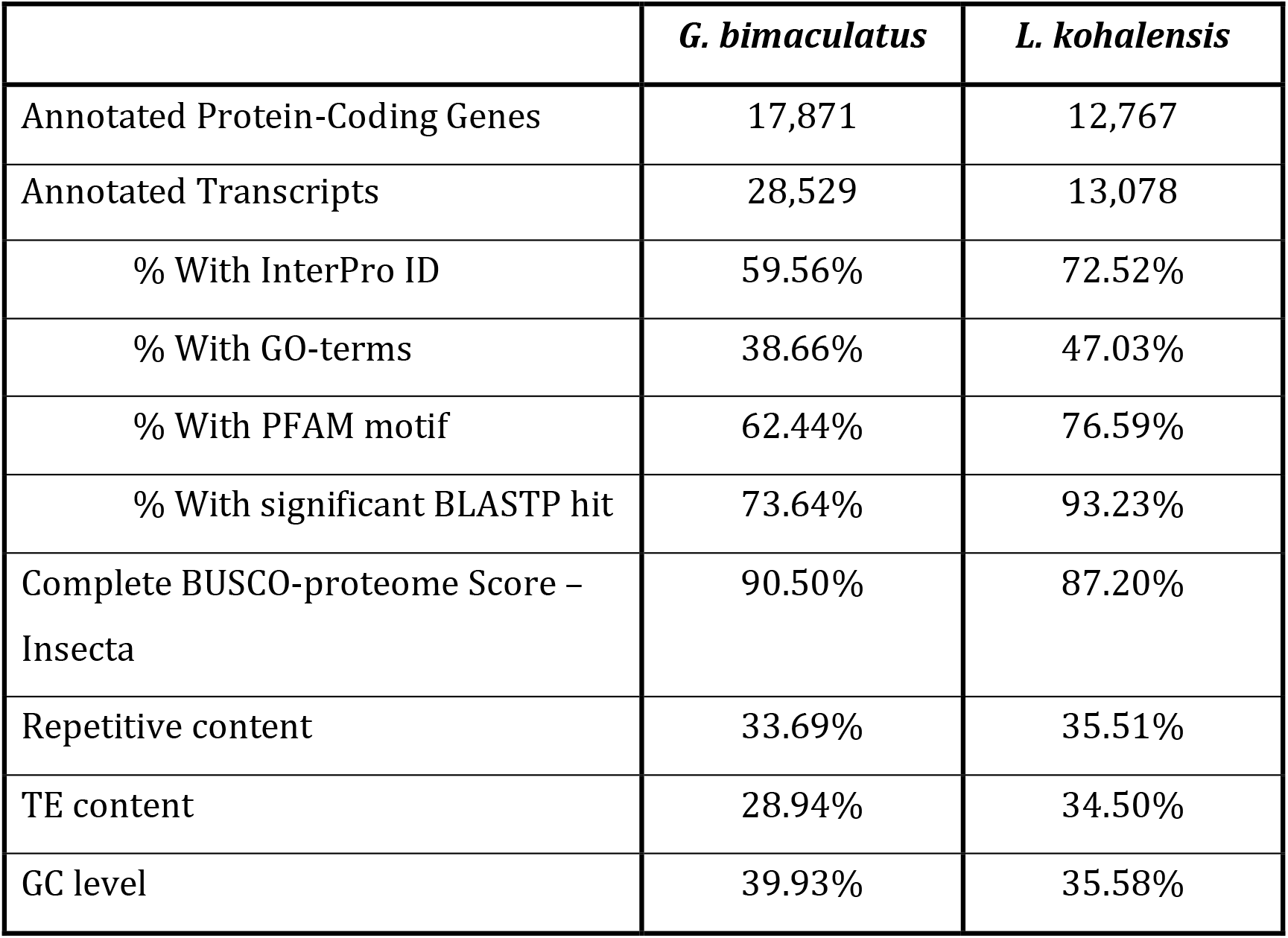
*Genome annotation summary for the crickets* G. bimaculatus *and* L. kohalensis

To obtain functional insights into the annotated genes, we ran InterProScan^19^ for all predicted protein sequences and retrieved their InterPro ID, PFAM domains, and Gene Ontology (GO) terms **(Table 2)**. In addition, we retrieved the best significant BLASTP hit (E-value < 1e-6) for 70-90% of the proteins. Taken together, these methods predicted functions for 75% and 94% of the proteins annotated for *G. bimaculatus* and *L. kohalensis* respectively. We created a novel graphic interface through which interested readers can access, search, BLAST and download the genome data and annotations (http://gbimaculatusgenome.rc.fas.harvard.edu).

### Abundant Repetitive DNA

We used RepeatMasker^20^ to determine the degree of repetitive content in the cricket genomes, using specific custom repeat libraries for each species. This approach identified 33.69% of the *G. bimaculatus* genome, and 35.51% of the *L. kohalensis* genome, as repetitive content **(Supplementary File 1)**. In *G. bimaculatus* the repetitive content density was similar throughout the genome, with the exception of scaffolds shorter than 1Mb (L90), which make up 10% of the genome and have a high density of repetitive content and low gene density **(Figure 1B)**. Because the repetitive content makes genome assemblies more challenging, as observed for the shortest scaffolds of *G. bimaculatus*, we cannot rule out the possibility that the lower contiguity of the *L. kohalensis* genome could lead us to underestimate its repetitive content. This caveat notwithstanding, we observed that transposable elements (TEs) accounted for 28.94% of the *G. bimaculatus* genome, and for 34.50% of the *L. kohalensis* genome. Although the overall proportion of genome made up of TEs was similar between the two cricket species, the proportion of each specific TE class varied greatly **(Figure 1C)**. In *L. kohalensis* the most abundant TE type was long interspersed elements (LINEs), accounting for 20.21% of the genome and 58.58% of the total TE content, while in *G. bimaculatus* LINEs made up only 8.88% of the genome and 30.68% of the total TE content. The specific LINE subtypes LINE1 and LINE3 appeared at a similar frequency in both cricket genomes (<0.5%), while the LINE2 subtype was over five times more represented in *L. kohalensis*, covering 10% of the genome (167 Mb). On the other hand, DNA transposons accounted for 8.61% of the *G. bimaculatus* genome, but only for 3.91% of the *L. kohalensis* genome.

### DNA methylation

CpG depletion, calculated as the ratio between observed versus the expected incidence of a cytosine followed by a guanine (CpG_o/e_), is considered a reliable indicator of DNA methylation. This is because spontaneous C to T mutations occur more frequently on methylated CpGs than unmethylated CpGs^16^. Thus, genomic regions that undergo methylation are eventually CpG-depleted. We calculated the CpG_o/e_ value for each predicted protein-coding gene for the two cricket species. In both species, we observed a clear bimodal distribution of CpG_o/e_ values (**Figure 2A**). One interpretation of this distribution is that the peak corresponding to lower CpG_o/e_ values contains genes that are typically methylated, and the peak of higher CpG_o/e_ contains genes that do not undergo DNA methylation. Under this interpretation, some genes have non-random differential DNA methylation in crickets. To quantify the genes in the two putative methylation categories, we set a CpG_o/e_ threshold as the value of the point of intersection between the two normal distributions (**Figure 2A**). After applying this cutoff, 44% of *G. bimaculatus* genes and 45% of *L. kohalensis* genes were identified as CpG-depleted.

A GO enrichment analysis of the genes above and below the CpG_o/e_ threshold defined above revealed clear differences in the predicted functions of genes belonging to each of the two categories. Strikingly, however, genes in each threshold category had functional similarities across the two cricket species (**Figure 3**). Genes with low CpG_o/e_ values, which are likely those undergoing methylation, were enriched for functions related to DNA replication and regulation of gene expression (including transcriptional, translational, and epigenetic regulation), while genes with high CpG_o/e_ values, suggesting little or no methylation, tended to have functions related to metabolism, catabolism, and sensory systems.

**Figure 3:**
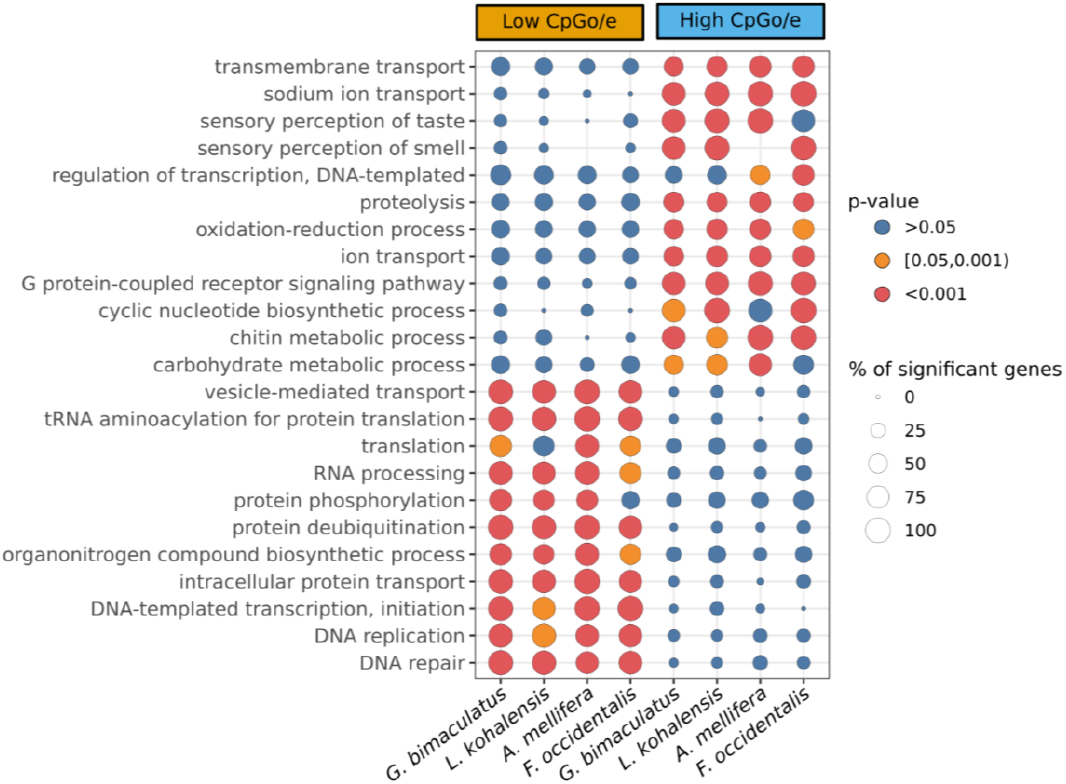
Functional analysis of high- and low-CpG_o/e_ genes: Enriched GO terms with a p-value<0.00001 in at least one of the eight categories, which are high CpG_o/e_ and low CpG_o/e_ genes of *G. bimaculatus, L. kohalensis, F. occidentalis*, and *A. mellifera.* The dot diameter is proportional to the percentage of significant genes with the GO term within the gene set. The dot color represents the p-value level: blue >0.05, orange [0.05, 0.001), red <0.001. Extended figure with all significant GO terms (p-value<0.05) available as **Supplementary Figure 3**.

To assess whether the predicted distinct functions of high- and low-CpG_o/e_ value genes were specific to crickets, or were a potentially more general trend of insects with DNA methylation systems, we analyzed the predicted functions of genes with different CpG_o/e_ values in the honeybee *Apis mellifera*, the first insect for which evidence for DNA methylation was robustly described and studied^21,22^, and the thrips *Frankliniella occidentalis*. We found that in both *F. occidentalis* and *A. mellifera*, CpG-depleted genes were enriched for similar functions as those observed in cricket CpG-depleted genes (**Figure 3 and Supplementary Figure 3**). Specifically, 23GO terms were significantly enriched in all four studied insects, and 15 additional GO terms were significantly enriched in the three hemimetabolous insects. In the same way, high CpG_o/e_ genes in all four insects were enriched for similar functions (8 GO-terms commonly enriched in all insects; **Supplementary Figure 3**).

Additionally, we observed that the proportion of species-specific genes was higher within the high CpG_o/e_ peak for all four insects (**Figure 2C**). In contrast, 86-96% of the genes belonging to the low CpG_o/e_ peak had an orthologous gene in at least one of the other studied insect species. Furthermore, we observed 2,182 orthogroups whose members always belonged to the low CpG_o/e_ peak in all four species, and 728 orthogroups whose members always belonged to the high CpG_o/e_ peak, indicating that orthologous genes are likely to share methylation state across these four insect species (**Figure 2B and Supplementary Figure 4**). Interestingly, 666 genes belonged to the low CpG_o/e_ peak in the three hemimetabolous species (*G. bimaculatus, L. kohalensis*, and *F. occidentallis)*, but to the high CpG_o/e_ peak in the holometabolous *A. mellifera*.

**Figure 4:**
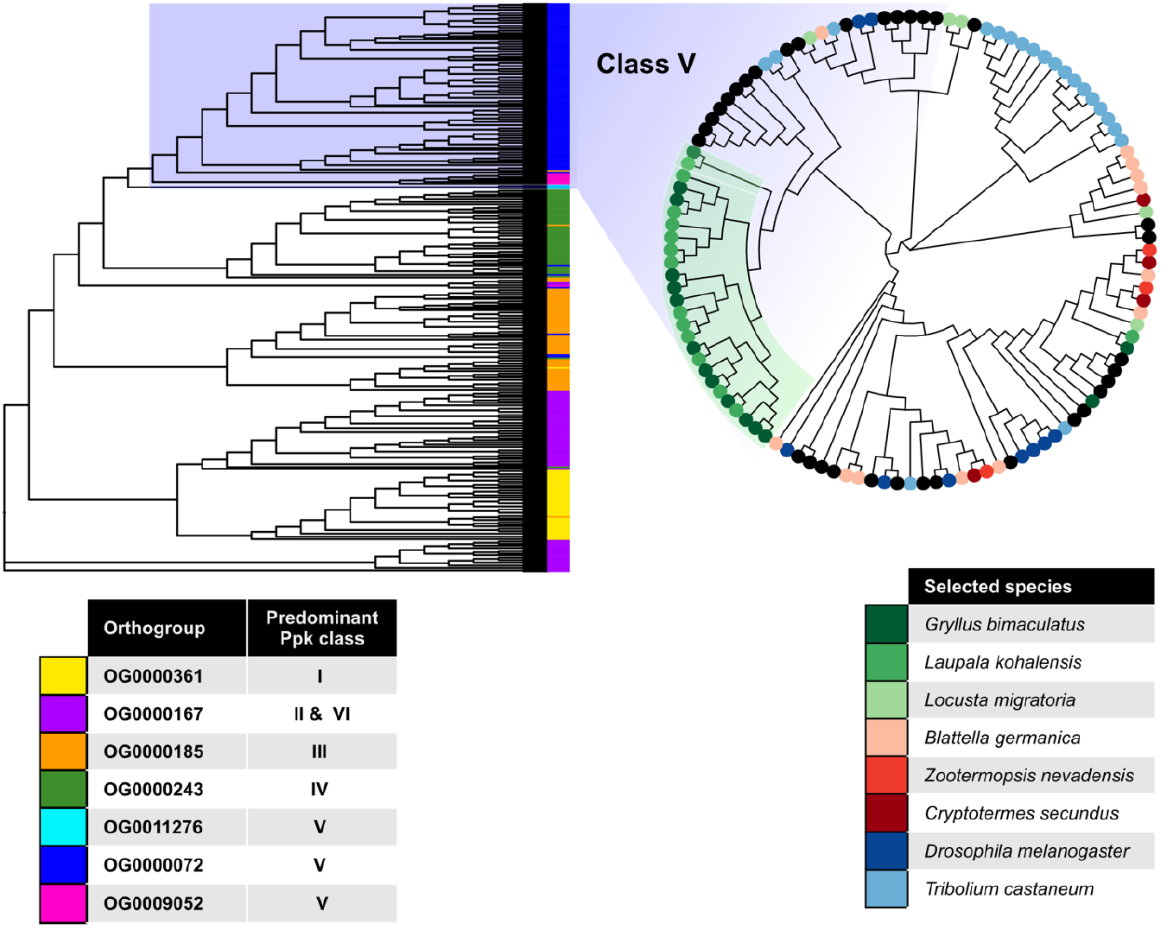
The *pickpocket* gene family class V is expanded in crickets. *pickpocket* gene tree with all the genes belonging to the seven OGs that contain the *D. melanogaster pickpocket* genes. All OGs predominantly contain members of a single *ppk* family. The OG0000167contains members of two *pickpocket* classes, II and VI. The orthogroup OG0000072 containing most *pickpocket* class V genes (circular cladogram) was significantly expanded in crickets relative to other insects.

**Figure 5:**
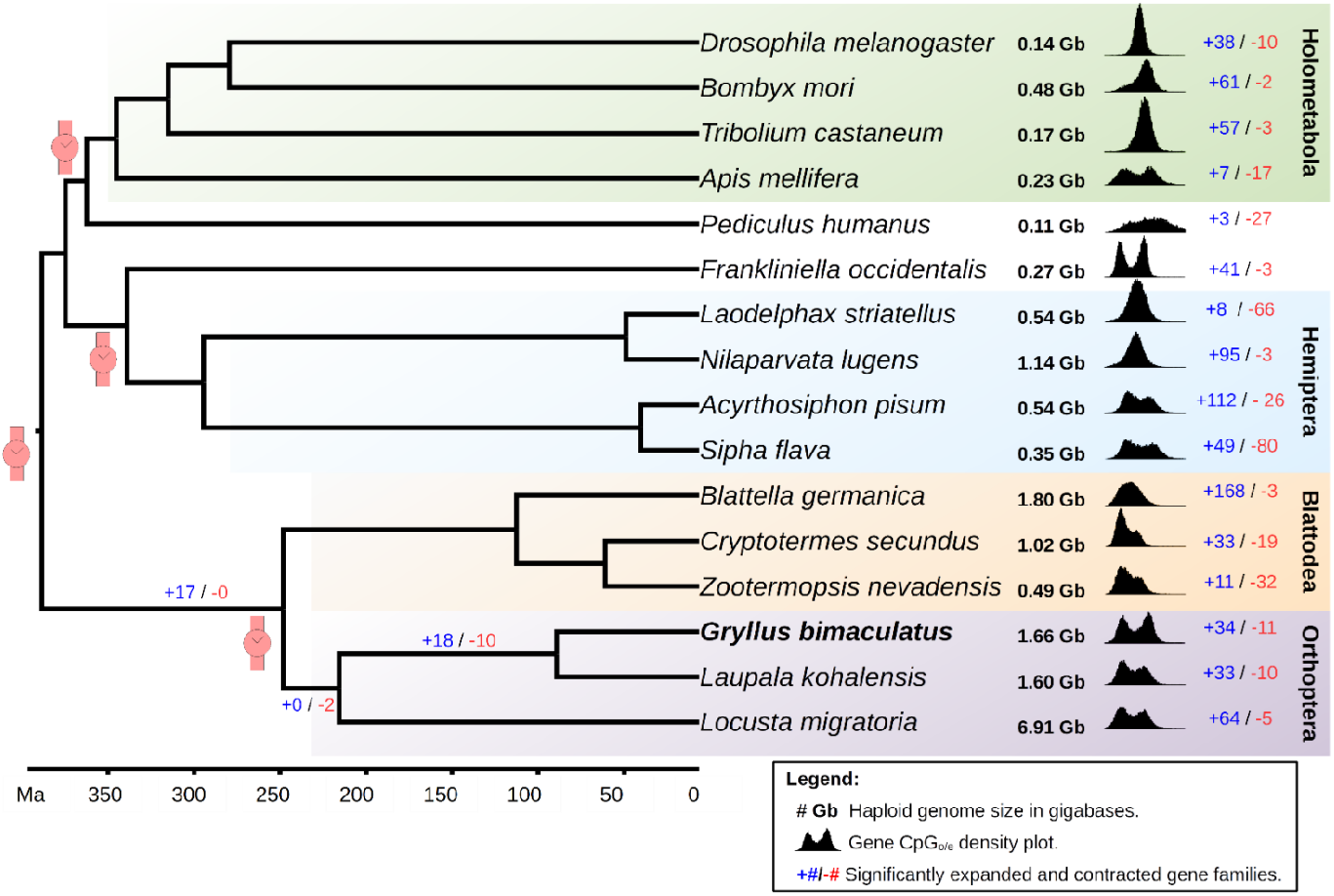
Cricket genomes in the context of insect evolution. A phylogenetic tree including 16 insect species calibrated at four different time points (red watch symbols) based on Misof, et al. ^23^, suggests that *G. bimaculatus* and *L. kohalensis* diverged ca. 89.2 Mya. The number of expanded (blue text) and contracted (red text) gene families is shown for each insect, and for the branches leading to crickets. The density plots show the CpG_o/e_ distribution for all genes for each species. The genome size in Gb was obtained from the genome fasta files (**Supplementary Table 1**).

Taken together, these results suggest that genes that are typically methylated tend to be more conserved across species, which could imply low evolutionary rates and strong selective pressure. To test this hypothesized relationship between low CpG_o/e_ and low evolutionary rates, we compared the dN/dS values of 1-to-1 orthologous genes belonging to the same CpG_o/e_ peak between the two cricket species. We found that CpG-depleted genes in both crickets had significantly lower dN/dS values than non-CpG-depleted genes (p-value<0.05; **Figure 2D**), consistent with stronger purifying selection on CpG-depleted genes.

### Phylogenetics and gene family expansions

To study the genome evolution of these cricket lineages, we compared the two cricket genomes with those of 14 additional insects, including members of all major insect lineages with special emphasis on hemimetabolous species. For each of these 16 insect genomes, we retrieved the longest protein per gene and grouped them into orthogroups (OGs), which we called “gene families” for the purpose of this analysis. The 732 OGs containing a single protein per insect, namely single copy orthologs, were used to infer a phylogenetic tree for these 16 species. The obtained species tree topology was in accordance with the currently understood insect phylogeny^23^. Then, we used the Misof et al. (2014) dated phylogeny to calibrate our tree on four different nodes, which allowed us to estimate that the two cricket species diverged circa 89.2 million years ago.

Our gene family expansion/contraction analysis using 59,516 OGs identified 18 gene families that were significantly expanded (p-value<0.01) in the lineage leading to crickets. In addition, we identified a further 34 and 33 gene family expansions specific to *G. bimaculatus* and *L. kohalensis* respectively. Functional analysis of these expanded gene families (**Supplementary File 2**) revealed that the cricket-specific gene family expansions included *pickpocket* genes, which are involved in mechanosensation in *Drosophila melanogaster* as described in the following section.

### Expansion of *pickpocket* genes

In *D. melanogaster*, the complete *pickpocket* gene repertoire is composed of 6 classes containing 31 genes. We found cricket orthologs of all 31 *pickpocket* genes across seven of our OGs, and each OG predominantly contained members of a single *pickpocket* class. We used all the genes belonging to these 7 OGs to build a *pickpocket* gene tree, using the predicted *pickpocket* orthologs from 16 insect species (**Figure 3**; **Supplementary Table 2**). This gene tree allowed us to classify the different *pickpocket* genes in each of the 16 species.

One orthogroup, which contained eight members of the *pickpocket* gene family of *D. melanogaster*, appeared to be significantly expanded to 14 or 15 members in crickets. Following the classification of *pickpocket* genes used in *Drosophila spp.^24^* we determined that the specific gene family expanded in crickets was *pickpocket* class V **(Figure 3).** In *D. melanogaster* this class contains eight genes: *ppk (ppk1), rpk (ppk2), ppk5, ppk8, ppk12, ppk17, ppk26*, and *ppk28* ^24^. Our analysis suggests that the class V gene family contains 15 and 14 genes in *G bimaculatus* and *L. kohalensis* respectively. In contrast, their closest analyzed relative, the locust *Locusta migratoria*, has only five such genes.

The *pickpocket* genes in crickets tended to be grouped in genomic clusters **(Figure 1B)**. For instance, in *G. bimaculatus* nine of the 15 class V *pickpocket* genes were clustered within a region of 900Kb, and four other genes appeared in two groups of two. In the *L. kohalensis* genome, although this genome is more fragmented than that of *G. bimaculatus* (**Supplementary Table 1**), we observed five clusters containing between two and five genes each.

In *D. melanogaster*, the *pickpocket* gene *ppk1* belongs to class V and is involved in functions related to stimulus perception and mechanotransduction^25^. For example, in larvae, this gene is required for mechanical nociception^26^, and for coordinating rhythmic locomotion^27^. *ppk* is expressed in sensory neurons that also express the male sexual behavior determiner *fruitless (fru)* ^28–30^.

To determine whether *pickpocket* genes in crickets are also expressed in the nervous system, we checked for evidence of expression of *pickpocket* genes in the publicly available RNA-seq libraries for the *G. bimaculatus* prothoracic ganglion^9^. This analysis detected expression (>20 transcripts per kilobase million, TPMs) of five *pickpocket* genes, four of them belonging to class V, in the *G. bimaculatus* nervous system. In the same ganglionic RNA-seq libraries, we also detected the expression of *fru* **(Supplementary Table 3**). Out of the four *pickpocket* genes, only one was detected in embryonic RNA-seq libraries. All four genes together with *fru* were detected in wild type leg transcriptomes, and their expression was found to be higher than wild type in a transcriptome from regenerating legs **(Supplementary Table 4)**.

## Discussion

### The importance of cricket genomes

Sequencing and analyzing genomes from underrepresented clades allow us to get a more complete picture of genome diversity across the tree of life, and can provide insights regarding their evolution. Since the first sequenced insect genome, that of *D. melanogaster*, was made publicly available in 2000^31^, the field of holometabolous genomics has flourished, and this clade became the main source of subsequent genomic information for insects. The first hemimetabolous genome was not available until ten years later, with the publication of the genome sequence and annotation of the Pea aphid *(Acyrthosiphon pisum)^32^.* When even more recently, polyneopteran genome sequences became available^33–36^, some of their distinct characteristics, such as their length and DNA methylation profiles, began to be appreciated. Genome data are also very important as they can help establish an animal species as tractable experimental models. *G. bimaculatus* is a common laboratory research animal used in neuroethology, developmental and regeneration biology studies^12,15^. It is our hope that the availability of the annotated genome presented here will encourage other researchers to adopt this cricket as a model organism, and facilitate development of new molecular genetic manipulation tools.

Moreover, we note that crickets are currently in focus as a source of animal protein for human consumption and for vertebrate livestock. Crickets possess high nutritional value, having a high proportion of protein for their body weight (>55%), and containing the essential linoleic acid as their most predominant fatty acid^37–39^. Specifically, the cricket *G. bimaculatus* has traditionally been consumed in different parts of the world including northeast Thailand, which recorded 20,000 insect farmers in 2011^40^. Studies have reported no evidence for toxicological effects related to oral consumption of *G. bimaculatus* by humans^41,42^, neither were genotoxic effects detected using three different mutagenicity tests^43^. A rare but known health risk associated with cricket consumption, however, is sensitivity and allergy to crickets^44,45^. Nevertheless, not only is the cricket *G. bimaculatus* considered generally safe for human consumption, several studies also suggest that introducing crickets into one’s diet may confer multiple health benefits ^46–48^.Crickets might therefore be part of the solution to the problem of feeding a worldwide growing population in a sustainable way. However, most of the crops and livestock that humans eat have been domesticated and subjected to strong artificial selection for hundreds or even thousands of years to improve their characteristics most desirable for humans, including size, growth rate, stress resistance, and organoleptic properties^49–52^. In contrast, to our knowledge, crickets have never been selected based on any food-related characteristic. The advent of genetic engineering techniques has accelerated domestication of some organisms^53^. These techniques have been used, for instance, to improve the nutritional value of different crops, or to make them tolerant to pests and climate stress^49,54^. Crickets are naturally nutritionally rich^39^, but in principle, their nutritional value could be further improved, for example by increasing vitamin content or Omega-3 fatty acids proportion. In addition, other issues that present challenges to cricket farming could potentially be addressed by targeted genome modification, which can be achieved in *G. bimaculatus* using Zinc finger nucleases, TALENs, or CRISPR/Cas9. These challenges include sensitivity to common insect viruses, aggressive behavior resulting in cannibalism, complex mating rituals, and relatively slow growth rate.

### Comparing cricket genomes to other insect genomes

The annotation of these two cricket genomes was done by combining *de novo* gene models, homology-based methods, and the available RNA-seq and ESTs. This pipeline allowed us to predict 17,871 genes in the *G. bimaculatus* genome, similar to the number of genes reported for other hemimetabolous insect genomes including the locust *L. migratoria* (17,307)^33^ and the termites *Cryptotermes secundus* (18,162)^55^, *Macrotermes natalensis* (16,140)^36^ and *Zootermopsis nevadensis*, (15,459)^35^. The slightly lower number of protein-coding genes annotated in *L. kohalensis* (12,767) may be due to the lesser amount of RNA-seq data available for this species, leading to higher assembly fragmentation, which challenges gene annotation. Nevertheless, the BUSCO scores are similar between the two crickets, and the proportion of annotated proteins with putative orthologous genes in other species (proteins with significant BLAST hits; see methods) for *L. kohalensis* is higher than for *G. bimaculatus*. This suggests the possibility that we may have successfully annotated most conserved genes, but that highly derived or species-specific genes might be missing from our annotations.

### TEs and genome size evolution

Approximately 35% of the genome of both crickets corresponds to repetitive content. This is substantially less than the 60% reported for the genome of *L. migratoria*^33^. This locust genome is one of the largest sequenced insect genomes to date (6.5 Gb) but has a very similar number of annotated genes (17,307) to those we report for crickets. We hypothesize that the large genome size difference between these orthopteran species is due to the TE content, which has also been correlated with genome size in multiple eukaryote species^56,57^.

Furthermore, we hypothesize that the differences in the TE composition between the two crickets are the result of abundant and independent TE activity since their divergence around 89.2 Mya. This, together with the absence of evidence for large genome duplication events in this lineage, leads us to hypothesize that the ancestral orthopteran genome was shorter than those of the crickets studied here (1.6 Gb for *G. bimaculatus* and 1.59 Gb for *L. kohalensis*) which are in the lowest range of orthopteran genome sizes^58^. In summary, we propose that the wide range of genome sizes within Orthoptera, reaching as high as 8.55 Gb in the locust *Schistocerca gregaria*, and 16.56 Gb in the grasshopper *Podisma pedestris^4,59^*, is likely due to TE activity since the time of the last orthopteran ancestor. These observations are consistent with the results reported by Palacios-Gimenez, et al.^60^ of massive and independent recent TE accumulation in four chromosome races of the grasshopper *Vandiemenella viatica*.

There is a clear tendency of polyneopteran genomes to be much longer than those of the holometabolous genomes **(Figure 4)**. Two currently competing hypotheses are that (1) the ancestral insect genome was small, and was expanded outside of Holometabola, and (2) the ancestral insect genome was large, and it was compressed in the Holometabola^3^. Our observations are consistent with the first of these hypotheses.

### DNA Methylation

Most holometabolan species, including well-studied insects like *D. melanogaster* and *Tribolium castaneum*, do not perform DNA methylation, or they do it at very low levels^6,61^. The honeybee *A. mellifera* was one of the first insects for which functional DNA methylation was described^21^. Although this DNA modification was initially proposed to be associated with the eusociality of these bees^22^, subsequent studies showed that DNA methylation is widespread and present in different insect lineages independently of social behavior ^5^. DNA methylation also occurs in other non-insect arthropods^62^.

While the precise role of DNA methylation in gene expression regulation remains unclear, our analysis suggests that cricket CpG-depleted genes (putatively hypermethylated genes) show signs of purifying selection, tend to have orthologs in other insects, and are involved in basic biological functions related to DNA replication and the regulation of gene expression. These enriched functions are in agreement with previous observations that DNA methylated genes in arthropods tend to perform housekeeping functions^6,63^. These predicted functions differ from those of the non-CpG depleted genes (putatively hypomethylated genes), which appear to be involved in signaling pathways, metabolism, and catabolism. These predicted functional categories may be conserved from crickets over circa 345 million years of evolution, as we also detect the same pattern in the honeybee and a thrips species.

Taken together, these observations suggest a potential relationship between DNA methylation, sequence conservation, and function for many cricket genes. Nevertheless, based on our data, we cannot determine whether the methylated genes are highly conserved because they are methylated, or because they perform basic functions that may be regulated by DNA methylation events. In the cockroach *Blattella germanica*, DNA methyltransferase enzymes and genes with low CpG_o/e_ values show an expression peak during the maternal to zygotic transition^64^, and functional analysis has shown that the DNA methyltransferase 1 is essential for early embryo development in this cockroach^65^. These results in cockroaches, together with our observations, leads us to speculate that at least in Polyneopteran species, DNA methylation might contribute to the maternal zygotic transition by regulating essential genes involved in DNA replication, transcription, and translation.

### *pickpocket* gene expansion

The *pickpocket* genes belong to the Degenerin/epithelial Na+ channel (DEG/ENaC) family, which were first identified in *Caenorhabditis elegans* as involved in mechanotransduction^25^. The same family of ion channels was later found in many multicellular animals, with a diverse range of functions related to mechanoreception and fluid–electrolyte homeostasis^66^.

Most of the information on their roles in insects comes from studies in *D. melanogaster.* In this fruit fly, *pickpocket* genes are involved in neural functions including NaCl taste^67^, pheromone detection ^68^, courtship behavior ^69^, and liquid clearance in the larval trachea^66^.

In *D. melanogaster* adults, the abdominal ganglia mediate courtship and postmating behaviors through neurons expressing *ppk* and *fru*^28–30^. In *D. melanogaster* larvae, *ppk* expression in dendritic neurons is required to control the coordination of rhythmic locomotion^27^. In crickets, the abdominal ganglia are responsible for determining song rhythm^70^. Moreover, we find that in *G. bimaculatus*, both *ppk* and *fru* gene expression are detectable in the adult prothoracic ganglion. These observations suggest the possibility that class V *pickpocket* genes could be involved in song rhythm determination in crickets through their expression in abdominal ganglia.

This possibility is consistent with the results of multiple quantitative trait locus (QTL) studies done in cricket species from the genus *Laupala*, which identified genomic regions associated with mating song rhythm variations and female acoustic preference^71^. The 179 scaffolds that the authors reported being within one logarithm of the odds (LOD) of the seven QTL peaks, contained five *pickpocket* genes, three of them from class V and two from class IV. One of the two class IV genes also appears within a QTL peak of a second experiment^7,72^. Xu and Shaw ^73^ found that a scaffold in a region of LOD score 1.5 of one of their minor linkage groups (LG3) contains *slowpoke*, a gene that affects song interpulse interval in *D. melanogaster*, and this scaffold also contains two class III pickpocket genes (**Supplementary Table 5**).

In summary, the roles of *pickpocket* genes in controlling rhythmic locomotion, courtship behavior, and pheromone detection in *D. melanogaster*, their appearance in genomic regions associated with song rhythm variation in *Laupala*, and their expression in *G. bimaculatus* abdominal ganglia, lead us to speculate that the expanded *pickpocket* gene family in cricket genomes could be playing a role in regulating rhythmic wing movements and sound perception, both of which are necessary for mating^15^. We note that Xu and Shaw ^73^ hypothesized that song production in crickets is likely to be regulated by ion channels, and that locomotion, neural modulation, and muscle development are all involved in singing^73^. However, further experiments, which could take advantage of the existing RNAi and genome modification protocols for *G. bimaculatus^13^*, will be required to test this hypothesis.

In conclusion, the *G. bimaculatus* genome assembly and annotation presented here is a source of information and an essential tool that we anticipate will enhance the status of this cricket as a modern functional genetics research model. This genome may also prove useful to the agricultural sector, and could allow improvement of cricket nutritional value, productivity, and reduction of allergen content. Annotating a second cricket genome, that of *L. kohalensis*, and comparing the two genomes, allowed us to unveil possible synapomorphies of cricket genomes, and to suggest potentially general evolutionary trends of insect genomes.

## Materials and Methods

### DNA isolation

The *G. bimaculatus* white-eyed mutant strain was reared at Tokushima University, at 29±1 °C and 30-50% humidity under a 10-h light, 14-h dark photoperiod. Testes of a single male adult of the *G. bimaculatus* white-eyed mutant strain were used for DNA isolation and shortread sequencing. We used DNA from testes of an additional single individual to make a long read PacBio sequencing library to close gaps in the genome assembly. Because sex differentiation in the cricket *G. bimaculatus* is determined by the XX/XO system^74^, genomic DNA extracted from males contains the full set of chromosomes; males were therefore chosen for genomic DNA isolation.

### Genome Assembly

Paired-end libraries were generated with insert sizes of 375 and 500 bp, and mate-pair libraries were generated with insert sizes of 3, 5, 10, and 20kb. Libraries were sequenced using the Illumina HiSeq 2000 and HiSeq 2500 sequencing platforms. This yielded a total of 127.4 Gb of short read paired-end data, that was subsequently assembled using the *de novo* assembler Platanus (v. 1.2.1)^75^. Scaffolding and gap closing were performed using total 138.2 Gb of mate-pair data. A further gap closing step was performed using long reads generated by the PacBio RS system. The 4.3 Gb of PacBio subread data were used to fill gaps in the assembly using PBjelly (v. 15.8.24)^76^.

### Repetitive Content Masking

We generated a custom repeat library for each of the two cricket genomes by combining the outputs from homology-based and *de novo* repeat identifiers, including the LTRdigest together with LTRharvest^77^, RepeatModeler/RepeatClassifier (www.repeatmasker.org/RepeatModeler), MITE tracker^78^, TransposonPSI (http://transposonpsi.sourceforge.net), and the databases SINEBase^79^ and RepBase^80^. We removed redundancies from the library by merging sequences that were greater than 80% similar with usearch ^81^, and classified them with RepeatClassifier. Sequences classified as “unknown” were searched with BLASTX against the 9,229 reviewed proteins of insects from UniProtKB/Swiss-Prot. Those sequences with a BLAST hit (E-value < 1e-10) against a protein not annotated as a transposase, transposable element, copia protein, or transposon were removed from the custom repeat library. The custom repeat library was provided to RepeatMasker version open-4.0.5 to generate the repetitive content reports, and to the MAKER2 pipeline to mask the genome.

### Protein-Coding Genes Annotation

We performed genome annotations through three iterations of the MAKER2 (v2.31.8) pipeline^18^ combining *ab-initio* gene models and evidence-based models. For the *G. bimaculatus* genome annotation, we provided the MAKER2 pipeline with the 43,595 *G. bimaculatus* nucleotide sequences from NCBI, an assembled developmental transcriptome ^82^, an assembled prothoracic ganglion transcriptome^9^, and a genome-guided transcriptome generated with StringTie^83^ using 30 RNA-seq libraries (accession numbers: DRA011174 and DDBJ DRA11117) mapped to the genome with HISAT2^84^. As alternative ESTs and protein sequences, we provided MAKER2 with 14,391 nucleotide sequences from *L. kohalensis* available at NCBI, and an insect protein database obtained from UniProtKB/Swiss-Prot^85^.

For the annotation of the *L. kohalensis* genome, we ran the MAKER2 pipeline with the 14,391 *L. kohalensis* nucleotide sequences from NCBI, the assembled *G. bimaculatus* developmental and prothoracic ganglion transcriptomes described above, and the 43,595 NCBI nucleotide sequences. As protein databases, we provided the insect proteins from UniProtKB/Swiss-Prot plus the proteins that we annotated in the *G. bimaculatus* genome.

For both crickets, we generated *ab-initio* gene models with GeneMark-ES^86^ in self-training mode, and with Augustus^87^ trained with BUSCO v3^17^. After each of the first two MAKER2 iterations, additional gene models were obtained with SNAP^88^ trained with the annotated genes.

Functional annotations were obtained using InterProScan^19^, which retrieved the InterProDomains, PFAM domains, and GO-terms. Additionally, we ran a series of BLAST rounds from more specific to more generic databases, to assign a descriptor to each transcript based on the best BLAST hit. The first round of BLAST was against the reviewed insect proteins from UniProtKB/Swiss-Prot. Proteins with no significant BLAST hits (E-value < 1e-6) went to a second round against all proteins from UniProtKB/TrEMBL, and those without a hit with E-value<1e-6 were used in the final round of BLAST against all proteins from UniProtKB/Swiss-Prot.

A detailed pipeline scheme is available in **Supplementary Figures 1 & 2,** and the annotation scripts are available on GitHub (https://github.com/guillemylla/Crickets_Genome_Annotation).

### Quality Assessment

Genome assembly statistics were obtained with assembly-stats (https://github.com/sanger-pathogens/assembly-stats). BUSCO (v3.1.0)^17^ was used to assess the level of completeness of the genome assemblies (‘-m geno’) as well as that of the gene annotations (‘-m prot’) at both arthropod (‘arthropoda_odb9’) and insect (‘insecta_odb9’) levels.

### CpG_o/e_ Analysis

We used the genome assemblies and their gene annotations from this study for the two cricket species, and retrieved publicly available annotated genomes from the other 14 insect species (**Supplementary Table 1**). The gene annotation files (in gff format) were used to obtain the amino-acid and CDS sequences for each annotated protein-coding gene per genome using gffread, with options “-y” and “-x” respectively. The CpG_o/e_ value per gene was computed as the observed frequency of CpGs (f_CpG_) divided by the product of C and G frequencies (f_c_ and f_G_) f_CpG_/f_C_*f_G_ in the longest CDS per gene for each of the 16 studied insects. CpG_o/e_ values larger than zero and smaller than two were retained and represented as density plots (**Figures 2 & 4**).

The distributions of gene CpG_o/e_ values per gene of the two crickets, the honeybee *A. mellifera*, and the thrips *F. occidentalis*, were fitted with a mixture of normal distributions using the mixtools R package^89^. This allowed us to obtain the mean of each distribution, the standard errors, and the interception point between the two distributions, which was used to categorize the genes into low CpG_o/e_ and high CpG_o/e_ bins. For these two bins of genes, we performed a GO-enrichment analysis (based on GO-terms previously obtained using InterProScan) of Biological Process terms using the TopGO package^90^ with all genes as universe, minimum node size of 10, the weight01 algorithm and the Fisher statistic. The GO terms with a p-value<0.05 were considered significantly enriched. Those GO terms significantly enriched in at least one gene set are shown in **Supplementary Figure 3**, and a subset of them with p-value<0.0001 are shown in **Figure 3**. In both figures, the size of the circle represents the percentage of enriched genes inside the set compared to all genes with the given GO term.

For each of the genes belonging to low and high CpG_o/e_ categories in each of the four insect species, we retrieved their orthogroup identifier from our gene family analysis, allowing us to assign putative methylation status to orthogroups in each insect. Then we used the UpSet R package^91^ to compute and display the number of orthogroups exclusive to each combination as an UpSet plot.

### dN/dS Analysis

We first aligned the longest predicted protein product of the single-copy-orthologs of all protein-coding genes between the two crickets (N=5,728) with MUSCLE (v3.8.31). Then, the amino-acid alignments were transformed into codon-based nucleotide alignments using the Pal2Nal software^92^. The resulting codon-based nucleotide alignments were used to calculate the pairwise dN/dS for each gene pair with the yn00 algorithm implemented in the PAML package^93^. Genes with dN or dS >2 were discarded from further analysis. The Wilcoxon-Mann-Whitney statistical test was used to compare the dN/dS values between genes with high and low CpG_o/e_ values in both insects.

### Gene Family Expansions and Contractions

Using custom Python scripts (see https://github.com/guillemylla/Crickets_Genome_Annotation) we obtained the longest predicted protein product per gene in each of the 16 studied insect species and grouped them into orthogroups (which we also refer to herein as “gene families”) using OrthoFinder v2.3.3^94^. The orthogroups (OGs) determined by OrthoFinder that contained a single gene per insect, namely putative one-to-one orthologs, were used for phylogenetic reconstruction. The proteins within each orthogroup were aligned with MUSCLE^95^ and the alignments trimmed with GBlocks (−t=p −b4=5 −b5=a)^96^. The trimmed alignments were concatenated into a single meta-alignment that was used to infer the species tree with FastTree2 (FastTreeMP-gamma)^97^.

To calibrate the species tree, we used the “chronos” function from the R package ape v5.3^98^, setting the common node between Blattodea and Orthoptera at 248 million years (my), the origin of Holometabola at 345 my, the common node between Hemiptera and Thysanoptera at 339 my, and the ancestor of hemimetabolous and holometabolous insects (root of the tree) at between 385 and 395 my. These time points were obtained from a phylogeny published that was calibrated with several fossils^23^.

The gene family expansion/contraction analysis was done with the CAFE software^99^. We ran CAFE using the calibrated species tree and the table generated by OrthoFinder with the number of genes belonging to each orthogroup in each insect. Following the CAFE manual, we first calculated the birth-death parameters with the orthogroups having less than 100 genes. We then corrected them by assembly quality and calculated the gene expansions and contractions for both large (>100 genes) and small (≤100) gene families. This allowed us to identify gene families that underwent a significant (p-value<0.01) gene family expansion or contraction on each branch of the tree. We proceeded to obtain functional information from those families expanded on our branches of interest (i.e. the origin of Orthoptera, the branch leading to crickets, and the branches specific to each cricket species.). To functionally annotate the orthogroups of interest, we first obtained the *D. melanogaster* identifiers of the proteins within each orthogroup, and retrieved the FlyBase Symbol and the FlyBase gene summary per gene using the FlyBase API^100^. Additionally, we ran InterProScan on all the proteins of each orthogroup and retrieved all PFAM motifs and the GO terms together with their descriptors. All of this information was summarized in tabulated files (**Supplementary File 2**), which we used to identify gene expansions with potentially relevant functions for insect evolution.

### *pickpocket* gene family expansion

Among the expanded gene families in crickets, we identified an orthogroup containing seven out of the eight *D. melanogaster pickpocket* class V genes, leading us to interpret that the *pickpocket* class V was significantly expanded in crickets. Subsequently, we retrieved the 6 additional orthogroups containing the completeset of *pickpocket* genes in *D. melanogaster*, and we assigned to each orthogroup the *pickpocket* class to which most of its *D. melanogaster* genes belonged according to Zelle and colleagues ^24^ (**Supplementary Table 2**). The protein sequences of all the members of the seven Pickpocket orthogroups were aligned with MUSCLE, and the *pickpocket* gene tree obtained with FastTree2 (FastTreeMP --gamma). The tips of the tree were colored based on the orthogroup to which they belong. A subset of the tree containing all the orthogroups that compose the entire *pickpocket* class V family was displayed as a circular cladogram (**Figure 3**), revealing an independent expansion of this family in *T. castaneum*.

To check for evidence of expression *pickpocket* genes in the cricket nervous system, we used the 21 RNA-seq libraries from prothoracic ganglion^9^ of *G. bimaculatus* available at NCBI GEO (PRJNA376023). Reads were mapped against the *G. bimaculatus* genome with RSEM^101^ using STAR^102^ as the mapping algorithm, and the number of expected counts and TPMs were retrieved for each gene in each library. The TPMs of the *pickpocket* genes and *fruitless* are shown in **Supplementary Table 3**. Genes with a sum of more than 20 TPMs across all samples were considered to be expressed in *G. bimaculatus* prothoracic ganglion. We further analyzed the *pickpocket* expression in the aggregated embryo RNA-seq dataset (DRA011174) and normal and regenerating legs RNA-seq dataset^103^ (DRR001985, DRR001986), using the same methodology.

## Acknowledgments

This work was supported by Harvard University and MEXT KAKENHI (No. 221S0002; 26292176; 17H03945). The computational infrastructure in the cloud used for the genome analysis was funded by AWS Cloud Credits for Research. The authors are grateful to Hiroo Saihara for his support in the management of a genome data server at Tokushima University.

## Author contributions statement

GY, SN, TM and CE designed experiments; TI and AT conducted sequencing by HiSeq and assembling short reads using the Platanus assembler; ST, YI, TW, MF and YM performed DNA isolation, gap closing of contigs and manual annotation; GY, TN, ST, TB and AAB conducted all other experiments and analyses; TM and CE funded the project; GY and CE wrote the paper with input from all authors.

## Data availability

The genome sequencing reads, RNA-seq reads, and the genome assembly for *Gryllus bimaculatus* were submitted to DDBJ and to NCBI under the accession number (PRJDB10609). The genome assembly and annotations can also be accessed and browsed at http://gbimaculatusgenome.rc.fas.harvard.edu.

## Code availability

The scripts used for genome annotation and analysis are available at GitHub (https://github.com/guillemylla/Crickets_Genome_Annotation).

## Supplementary Materials

These Supplementary Materials consist of the following:

**Supplementary Figure 1:**
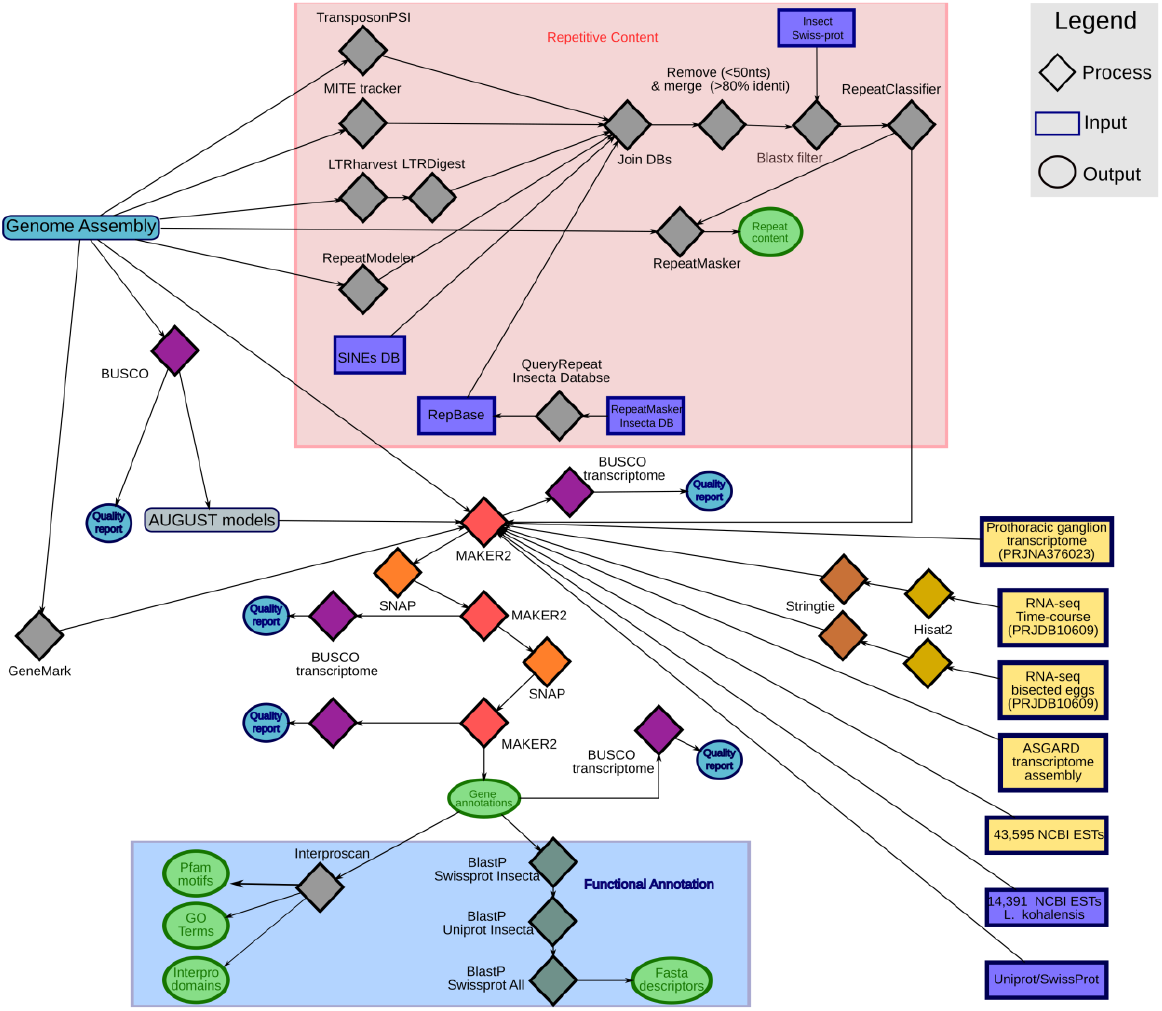
Schematic of *G. bimaculatus* genome annotation pipeline. Rectangles represent data inputs: yellow rectangles represent *G. bimaculatus* data; purple rectangles represent data from other species or databases. Diamonds represent computational processes: gray diamonds indicate processes executed a single time; non-gray diamonds of the same color indicate the same process. Circles indicate outputs: blue circles indicate quality controls; green circles indicate annotations. Scripts available at GitHub https://github.com/guillemylla/Crickets_Genome_Annotation.

**Supplementary Figure 2:**
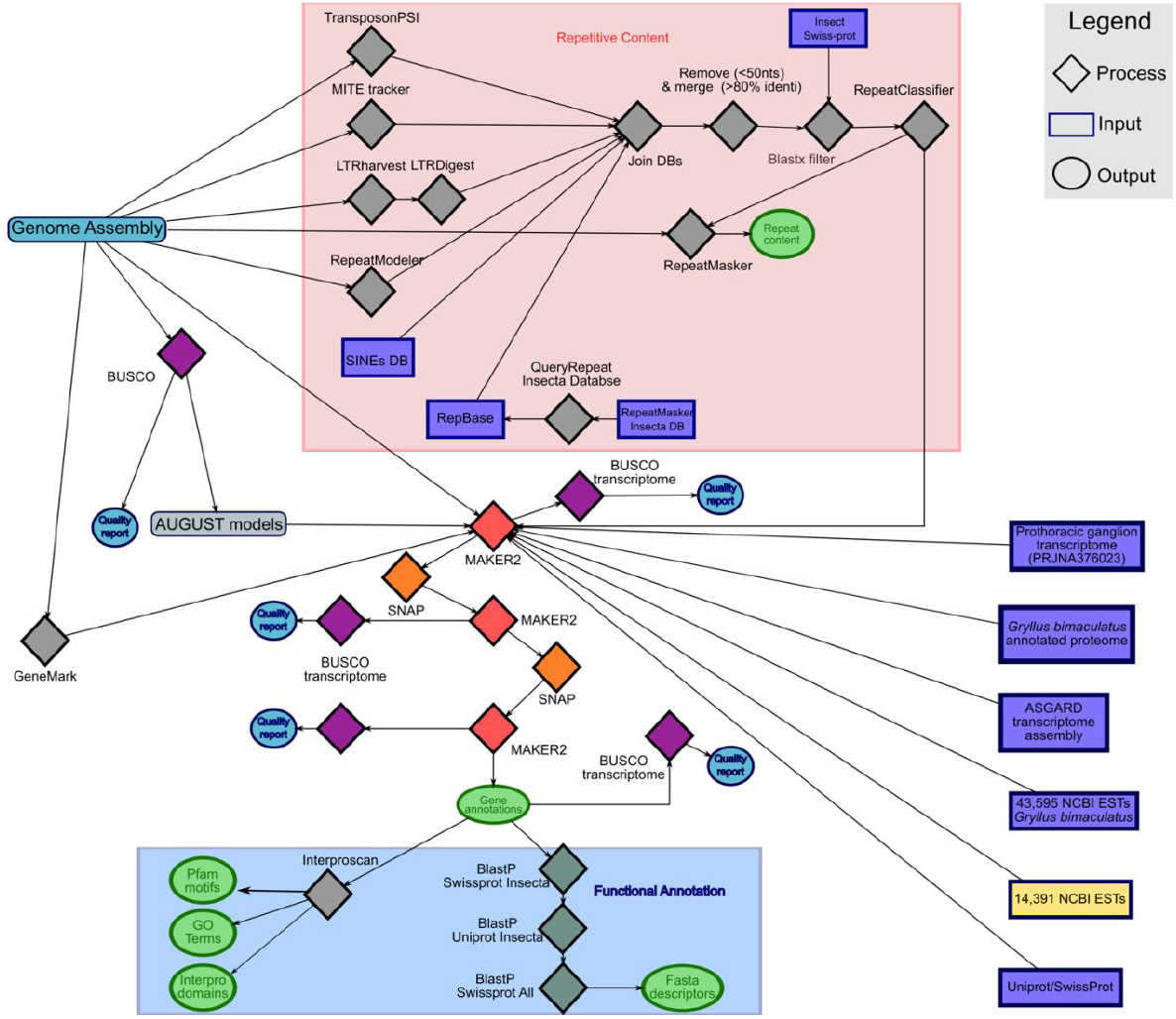
Scheme of *L. kohalensis* genome annotation pipeline. All symbols as per **Supplementary Figure 1**.

**Supplementary Figure 3:**
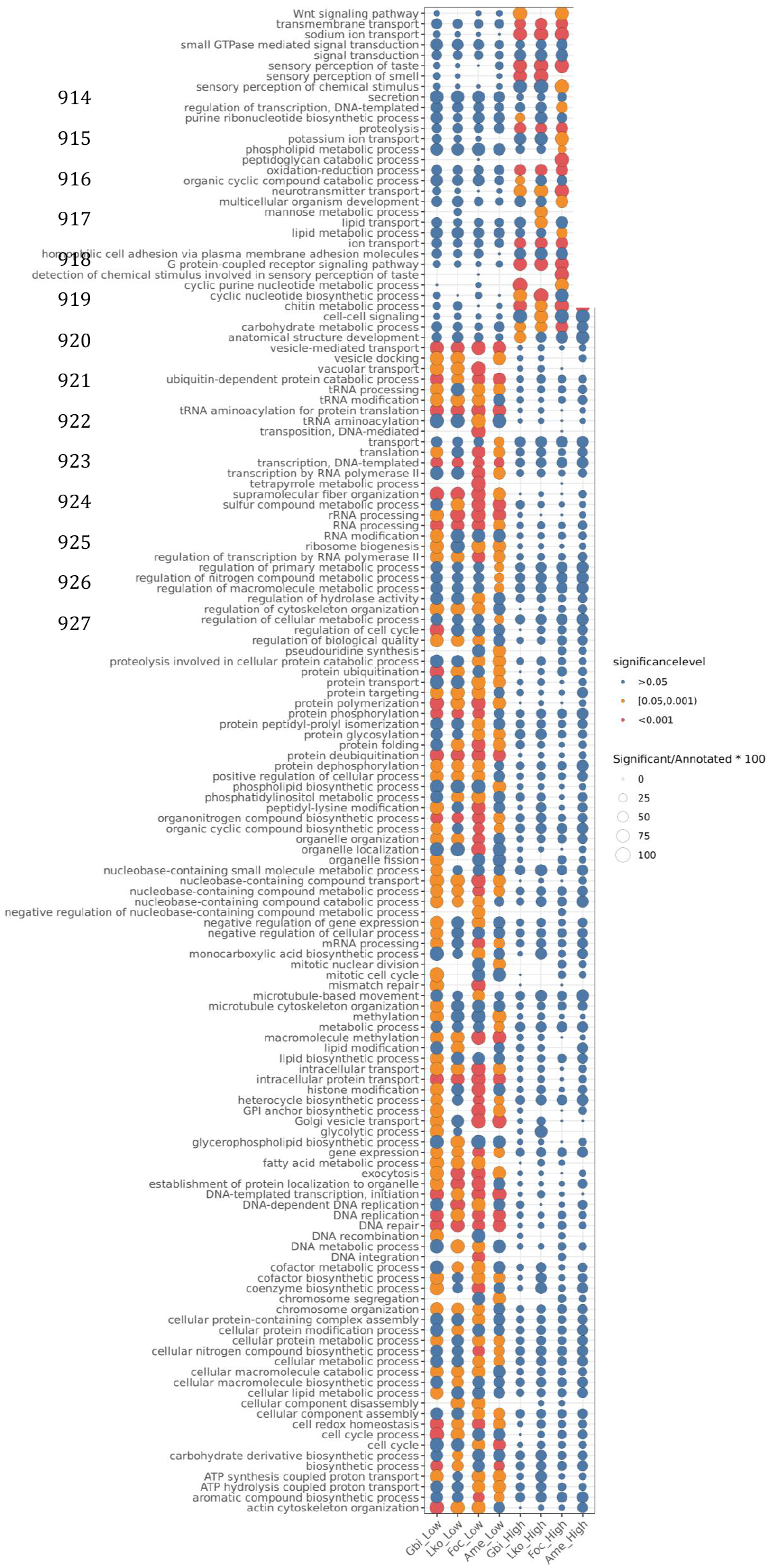
Enriched GO-terms among genes with high or low CpG_o/e_ levels. This plot shows the enriched GO terms with p-value<0.05 in at least one of the eight categories which are the high CpG_o/e_ and low CpG_o/e_ genes of *G. bimaculatus* (Gbi), *L. kohalensis* (Lko), *F. occidentalis* (Foc), and *A. mellifera* (Ame). The dot diameter is proportional the percentage of significant genes with the GO term within the gene set. The dot color represents the p-value level: blue >0.05, orange [0.05, 0.001), red <0.001.

**Supplementary Figure 4:**
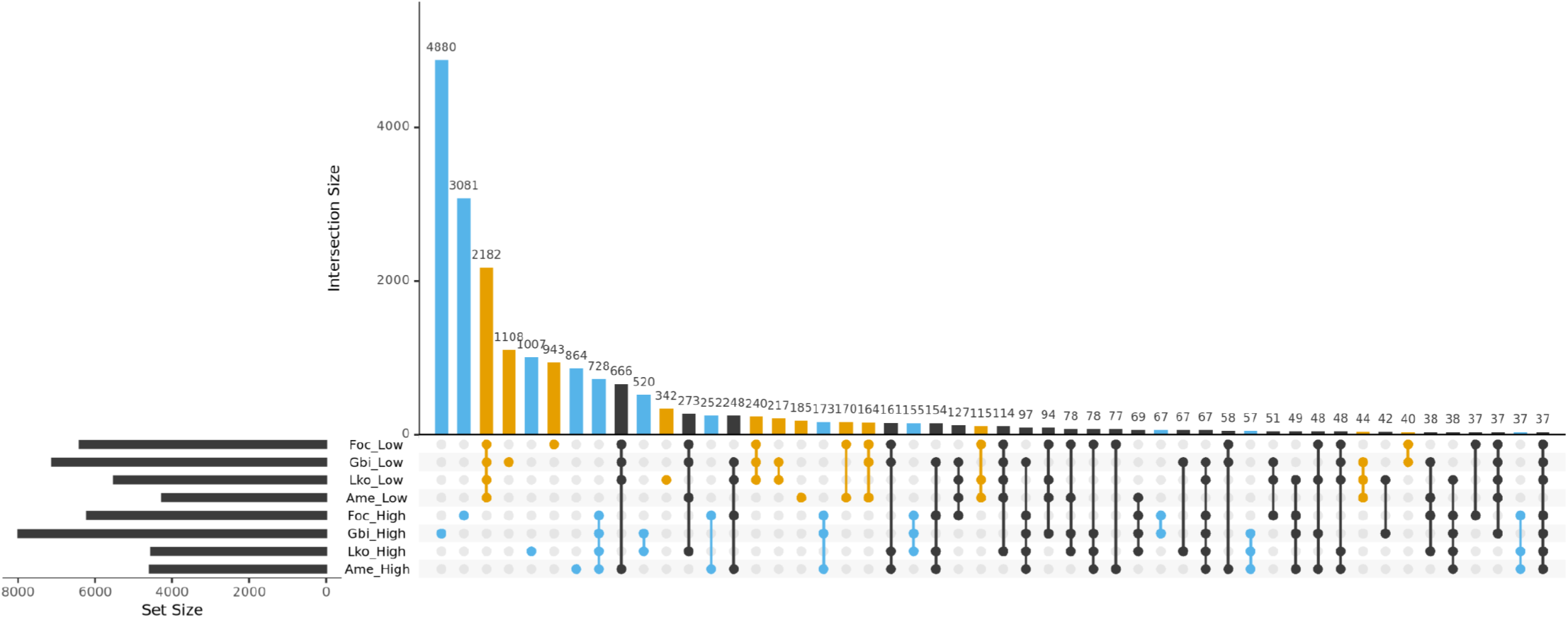
UpSet plot of orthologous genes within the high and low CpG_o/e_ value categories. Top 50 intersections of orthogroups (OGs) that are common across the eight different categories, which are the high CpG_o/e_ and low CpG_o/e_ genes for *G. bimaculatus* (Gbi), *L. kohalensis* (Lko), *F. Occidentalis* (Foc), and *A. mellifera* (Ame). Blue color indicates OGs that contain genes that only belong to the high CpG_o/e_ peak, and yellow OGs contain genes that only belong to the low CpG_o/e_ peak.

**Supplementary File 1:**
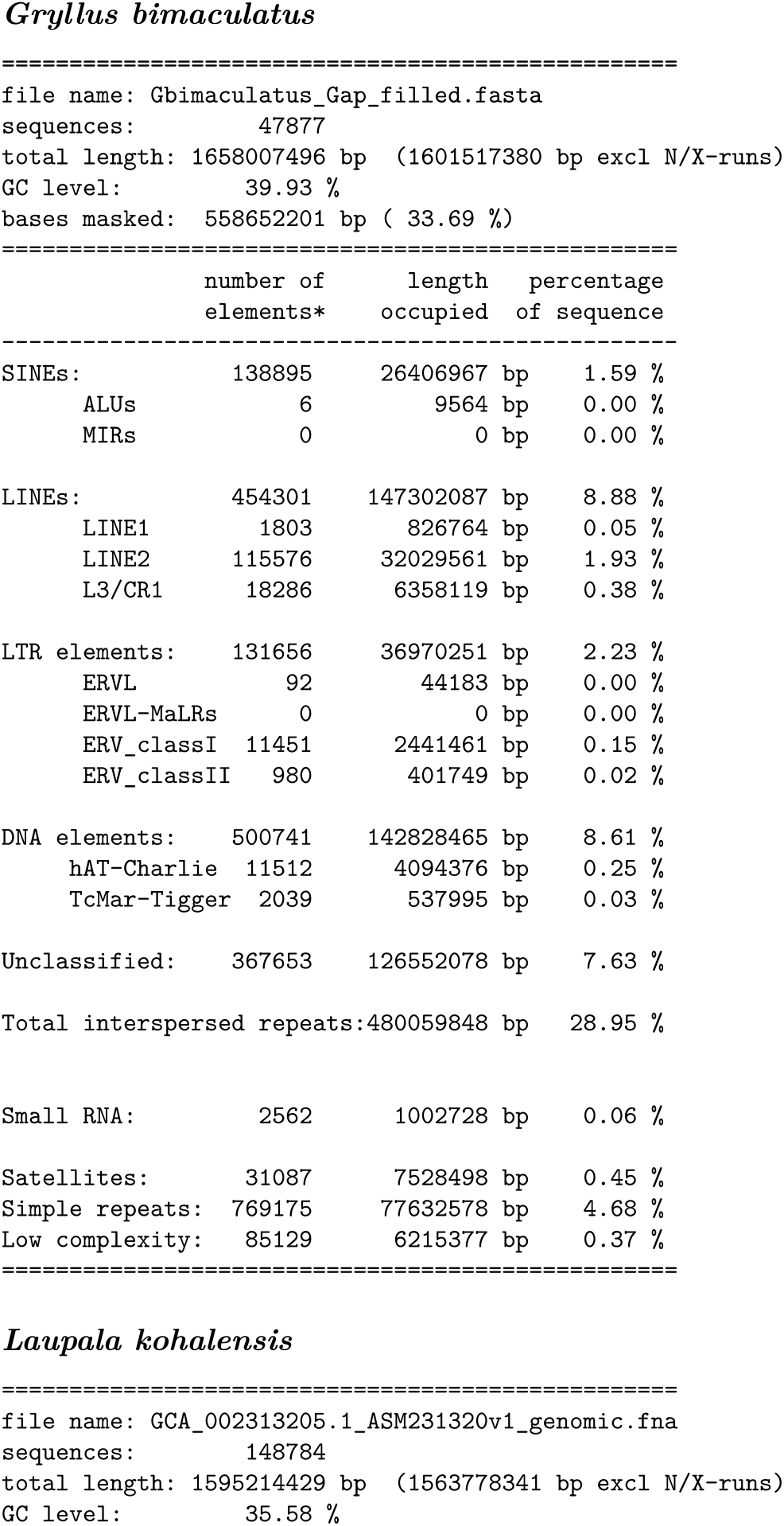
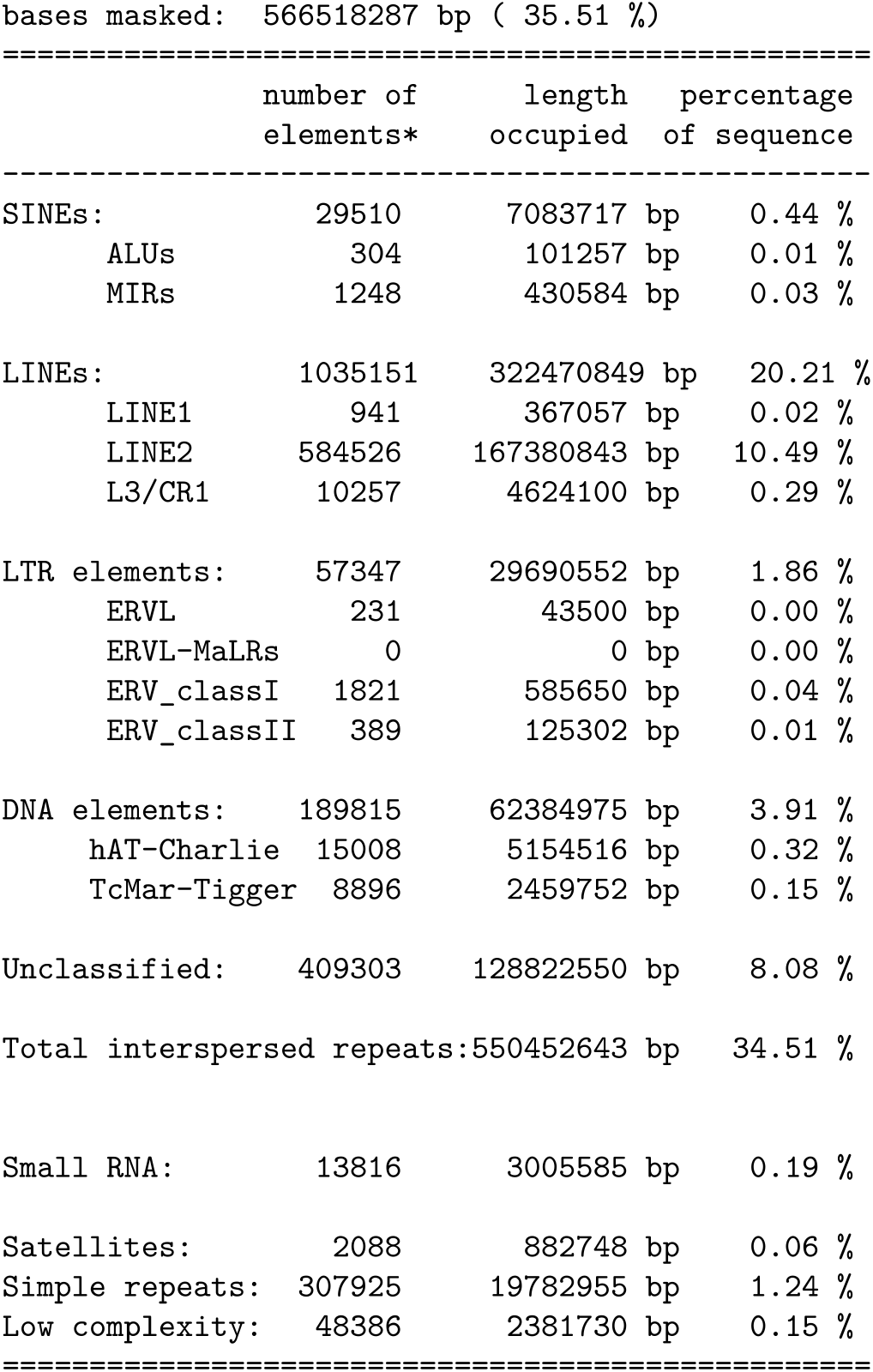
RepeatMasker summaries. Report of the repeat content in the genomes of *G. bimaculatus* and *L. kohalensis* generated by RepeatMasker using custom libraries.

**Supplementary File 2: Gene family expansions in crickets.** Gene families (Orthogroups) significantly expanded in the lineage leading to crickets (tab 1), expanded in *G. bimaculatus* (tab 2), and expanded in *L. kohalensis* (tab 3). For each expanded orthogroup (OG), we report the expansion size as the number of genes gained, and the functional information about the OG. The functional information consists of the list of PFAMs and GO terms associated with the genes within the OG, and the list of *D. melanogaster* genes within the OG with their FlyBase summaries.

*See file “Supplementary_File_2_GeneExpansions.xls”*

**Supplementary Table 1: Genome assembly information for the 16 insect genomes analyzed.** For each genome, we show the database that the assembly was retrieved from, the assembly file name, the accession code, the assembly statistics obtained with assembly-stats software (https://github.com/sanger-pathogens/assembly-stats) and the BUSCO v3.1.0 reports at Arthropoda and Insecta levels.

*See file “Supplementary_Table_1_GenomeStats.xls”*

**Supplementary Table 2:**
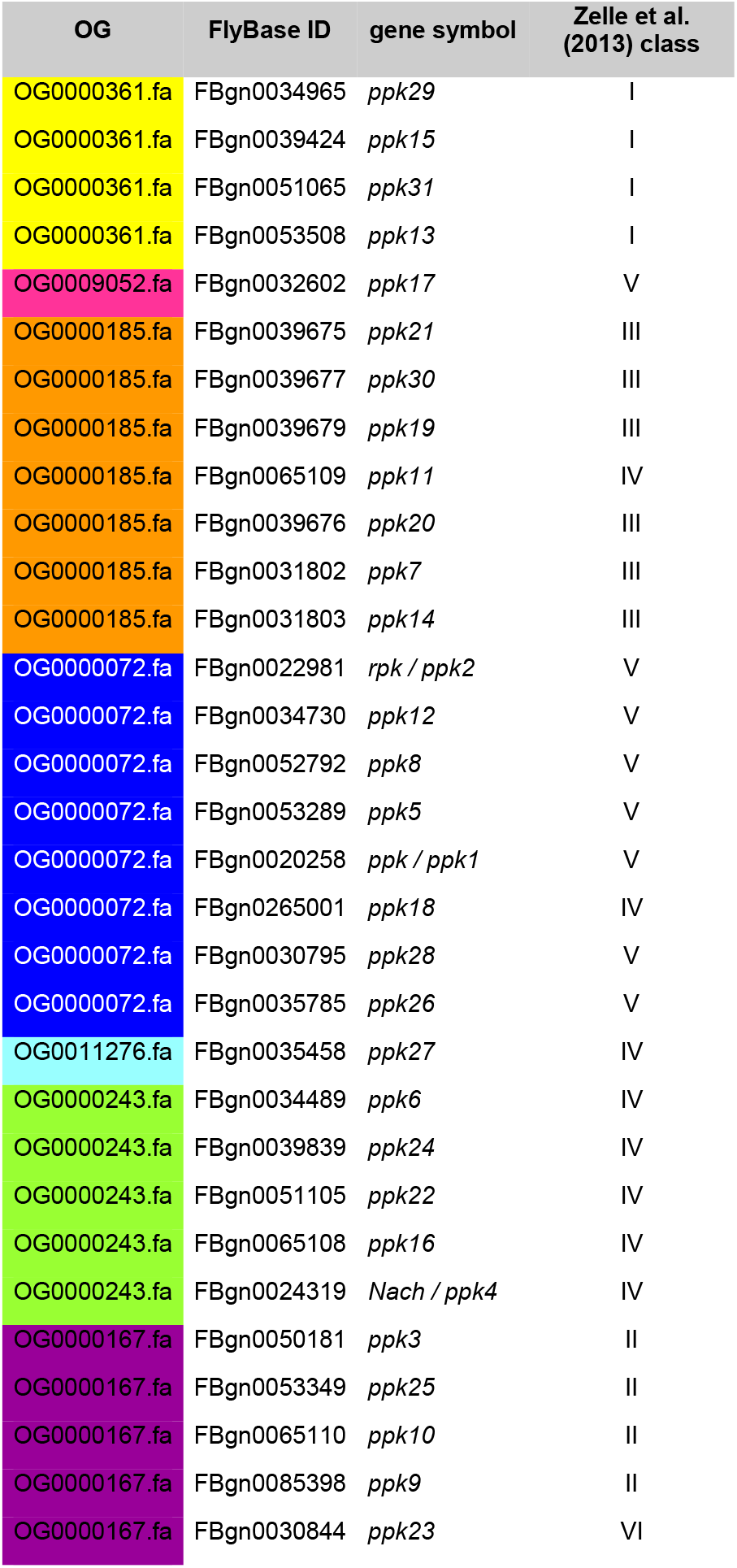
The orthogroups (OG) containing the 31 *D. melanogaster pickpocket* genes, with their FlyBase ID, symbol, and class according to Zelle et al. (2013).

| OG | FlyBase ID | gene symbol | Zelle et al.<br>(2013) class |
| --- | --- | --- | --- |
| OG0000361.fa | FBgn0034965 | *ppk29* | I |
| OG0000361.fa | FBgn0039424 | *ppk15* | I |
| OG0000361.fa | FBgn0051065 | *ppk31* | I |
| OG0000361.fa | FBgn0053508 | *ppk13* | I |
| OG0009052.fa | FBgn0032602 | *ppk17* | V |
| OG0000185.fa | FBgn0039675 | *ppk21* | III |
| OG0000185.fa | FBgn0039677 | *ppk30* | III |
| OG0000185.fa | FBgn0039679 | *ppk19* | III |
| OG0000185.fa | FBgn0065109 | *ppk11* | IV |
| OG0000185.fa | FBgn0039676 | *ppk20* | III |
| OG0000185.fa | FBgn0031802 | *ppk7* | III |
| OG0000185.fa | FBgn0031803 | *ppk14* | III |
| OG0000072.fa | FBgn0022981 | *rpk / ppk2* | V |
| OG0000072.fa | FBgn0034730 | *ppk12* | V |
| OG0000072.fa | FBgn0052792 | *ppk8* | V |
| OG0000072.fa | FBgn0053289 | *ppk5* | V |
| OG0000072.fa | FBgn0020258 | *ppk / ppk1* | V |
| OG0000072.fa | FBgn0265001 | *ppk18* | IV |
| OG0000072.fa | FBgn0030795 | *ppk28* | V |
| OG0000072.fa | FBgn0035785 | *ppk26* | V |
| OG0011276.fa | FBgn0035458 | *ppk27* | IV |
| OG0000243.fa | FBgn0034489 | *ppk6* | IV |
| OG0000243.fa | FBgn0039839 | *ppk24* | IV |
| OG0000243.fa | FBgn0051105 | *ppk22* | IV |
| OG0000243.fa | FBgn0065108 | *ppk16* | IV |
| OG0000243.fa | FBgn0024319 | *Nach / ppk4* | IV |
| OG0000167.fa | FBgn0050181 | *ppk3* | II |
| OG0000167.fa | FBgn0053349 | *ppk25* | II |
| OG0000167.fa | FBgn0065110 | *ppk10* | II |
| OG0000167.fa | FBgn0085398 | *ppk9* | II |
| OG0000167.fa | FBgn0030844 | *ppk23* | VI |

**Supplementary Table 3: *pickpocket* gene expression levels in the *G. bimaculatus* prothoracic ganglion.** Expression in TPMs of *fruitless* and *pickpocket* genes in each RNA-seq library generated from adult male prothoracic ganglia previously generated by Fisher and colleagues (2018). Genes with read sum across samples > 20 TPMs across samples are highlighted.

**Supplementary Table 4: *pickpocket* gene expression levels in the *G. bimaculatus* embryo and regenerating legs.** Expression in TPMs of *fruitless* and *pickpocket* genes in the aggregated embryo RNA-seq dataset, control legs and regenerating legs. Genes with read sum across samples > 20 TPMs across samples in the prothoracic ganglion (Supplementary Table 3) are highlighted.

See file “Supplementary_Table_3-4.xls”

**Supplementary Table 5:** *pickpocket* genes present in previous QTL analyses examining the genetic basis for sound-based cricket courtship behavior variation. Genomic position information for the *L. kohalensis pickpocket* genes found in linkage groups (LG) in previously published QTL analyses (Blankers, Oh & Shaw 2018; Shaw & Lesnick, 2009) examining mating song rhythm variations and female acoustic preference in the genus *Laupala.*

|  |  |  |  |  |  |  |  | Table S3 and S6<br>(Blankers, Oh, & Shaw, 2018) |  | Table S4 (Blankers,<br>Oh, Bombarely, & Shaw, 2018) | Table 2 (Xu<br>and Shaw, 2019) |
| --- | --- | --- | --- | --- | --- | --- | --- | --- | --- | --- | --- |
| Scaff names Shaw | Scaff Names<br>NCBI | start | end | width | strand | Name | Ppk class | LG | proximity | LG | LG |
| Lko057S000409 | NNCF01126148.1 | 1083057 | 1116038 | 32982 | + | Lko_01144 | Class IV | 1 | LOD1 | 1 |  |
| Lko057S000550 | NNCF01126289.1 | 666338 | 667949 | 1612 | - | Lko_06470 | Class IV | 3 | LOD2 |  |  |
| Lko057S005538 | NNCF01131273.1 | 20948 | 31450 | 10503 | - | Lko_31867 | Class V | 4 | LOD1 |  |  |
| Lko057S005538 | NNCF01131273.1 | 6676 | 8154 | 1479 | - | Lko_31866 | Class V | 4 | LOD1 |  |  |
| Lko057S005538 | NNCF01131273.1 | 43198 | 60736 | 17539 | - | Lko_31869 | Class V | 4 | LOD1 |  |  |
| Lko057S000206 | NNCF01125945.1 | 353321 | 357106 | 3786 | - | Lko_06341 | Class III |  |  |  | 3 |
| Lko057S000206 | NNCF01125945.1 | 404113 | 432386 | 28274 | - | Lko_06342 | Class III |  |  |  | 3 |

